# Lineage-mosaic and mutation-patched spike proteins for broad-spectrum COVID-19 vaccine

**DOI:** 10.1101/2022.01.25.477789

**Authors:** Yangtao Wu, Shaojuan Wang, Yali Zhang, Lunzhi Yuan, Qingbing Zheng, Min Wei, Yang Shi, Zikang Wang, Jian Ma, Kai Wang, Meifeng Nie, Jin Xiao, Zehong Huang, Peiwen Chen, Huilin Guo, Miaolin Lan, Jingjing Xu, Wangheng Hou, Yunda Hong, Dabing Chen, Hui Sun, Hualong Xiong, Ming Zhou, Che Liu, Wenjie Guo, Huiyu Guo, Jiahua Gao, Zhixiong Li, Haitao Zhang, Xinrui Wang, Shaowei Li, Tong Cheng, Qinjian Zhao, Yixin Chen, Ting Wu, Tianying Zhang, Jun Zhang, Hua Cao, Huachen Zhu, Quan Yuan, Yi Guan, Ningshao Xia

**Author notes:** These authors contributed equally to this work (Yangtao Wu, Shaojuan Wang, Yali Zhang, Lunzhi Yuan, Qingbing Zheng, Min Wei, Yang Shi). Corresponding author. (T. Z.), (J. Z.), (H. Z.), (H. C.), (Q. Y.), (Y. G.), (N. X.).

## Abstract

The widespread SARS-CoV-2 in humans results in the continuous emergence of new variants. Recently emerged Omicron variant with multiple spike mutations sharply increases the risk of breakthrough infection or reinfection, highlighting the urgent need for new vaccines with broad-spectrum antigenic coverage. Using inter-lineage chimera and mutation patch strategies, we engineered a recombinant monomeric spike variant (STFK1628x), which showed high immunogenicity and mutually complementary antigenicity to its prototypic form (STFK). In hamsters, a bivalent vaccine comprised of STFK and STFK1628x elicited high titers of broad-spectrum antibodies to neutralize all 14 circulating SARS-CoV-2 variants, including Omicron; and fully protected vaccinees from intranasal SARS-CoV-2 challenges of either the ancestral strain or immune-evasive Beta variant. Strikingly, the vaccination of hamsters with the bivalent vaccine completely blocked the within-cage virus transmission to unvaccinated sentinels, for either the ancestral SARS-CoV-2 or Beta variant. Thus, our study provides new insights and antigen candidates for developing next-generation COVID-19 vaccines.

## Introduction

The ongoing coronavirus disease 2019 (COVID-19) pandemic caused by the severe acute respiratory syndrome coronavirus 2 (SARS-CoV-2) poses an unbearable public health burden. The SARS-CoV-2 spike mainly contains three immunogenic domains as targets of neutralizing antibody (nAb), the N-terminal domain (NTD), the receptor-binding domain (RBD), and the subunit 2 (S2), thereby serving as the essential antigen of COVID-19 vaccines. Though several COVID-19 vaccines are available, the constant emergence of SARS-CoV-2 variants is challenging against the protective efficacy of vaccination. Viral genome mutations may alter the biological phenotypes of SARS-CoV-2 in many aspects, such as viral infectivity, pathogenicity, and antigenicity. Critically, the amino-acid substitutions in the antigenic sites of the spike protein may enable viruses to escape from naturally acquired and vaccine-induced immunity (*1*). Among the variants currently identified as variants of concern (VOCs) or variants of interest (VOIs), many were able to cause immune escape. The Beta (B.1.351) variant, first identified in South Africa, was found to cause a 6.5-8.6-fold decrease in nAb titers raised by existing mRNA vaccines (*2*). Besides, the Gamma (P.1), Delta (B.1.617.2), and Mu (B.1.621) variants also caused a 3.8-4.8-, 2.9- and 9.1-fold nAb decrease, respectively, according to previous reports (*3, 4*). The recently emerged Omicron (B.1.1.529) variant, being firstly identified in November 2021 and rapidly designated as a new VOC, has shown elevated risks in causing risk of breakthrough infection or reinfection due to its multiple spike mutations altering viral antigenicity (*5-9*).

Most of the currently licensed COVID-19 vaccines were designed based on the SARS-CoV-2 prototype spike; their vaccine effectiveness (VE) appeared to be compromised in countering variants with immune evasion. The phase 3 clinical trial of AZD1222 indicated that this vaccine was ineffective against mild-to-moderate COVID-19 disease due to the Beta variant (*10*). For Omicron, a recent study showed that the efficacy of two doses of BNT162b2 against symptomatic illnesses caused by this variant was only about 30%, whereas AZD1222 did not show a significant protective effect (*11*). Several vaccine manufacturers have announced their plans of antigen updates in developing vaccines with broad-spectrum protection (*12, 13*). An ideal goal is to develop a new antigen providing a broad-spectrum coverage for all SARS-CoV-2 variants that resist nAbs raised by the prototypic spike. However, it remains hugely challenging to achieve this goal.

In this study, using inter-lineage chimera and mutation patch strategies, we generated a serial of monomeric spike ectodomain proteins harboring multi-site mutations from different VOC/VOI variants. Our evaluations demonstrated a chimeric spike protein of STFK1628x, containing NTD from B.1.620 lineage, RBD-S2 from the Gamma variant, and additional RBD mutation patches from the Delta variant, showed mutually complementary antigenicity to the ancestral spike-derived STFK protein. The bivalent vaccine of STFK plus STFK1628x exhibited high immunogenicity to elicit high titers of broad-spectrum neutralizing antibodies to protect against *in vivo* challenges with the ancestral SARS-CoV-2 and Beta variant in hamsters. More importantly, we further evidenced the new bivalent vaccine could block the within-cage virus transmission from vaccinated hamsters to unvaccinated sentinels. Overall, our findings shed light on the understandings of antigenic and immunogenic characteristics of SARS-CoV-2 spike variants, also providing antigen candidates for developing next-generation COVID-19 vaccines.

## Results

### Monomeric spike ectodomain STFK protein is highly immunogenic in rodents and nonhuman primates

Several studies had demonstrated the immunogenicity of recombinant SARS-CoV-2 spike ectodomain protein in trimeric forms (*14-17*). However, introducing the exogenous trimerization domain may elicit an unexpected immune response(*14*). Moreover, the recombinant trimer protein may dissociate during the cell cultures and downstream purification process, which decreases the yield of homogeneous trimeric proteins for vaccine production. Therefore, we tried to design and produce monomeric spike proteins for the COVID-19 vaccine to address these issues. Although numerous high-resolution structures of SARS-CoV-2 spike trimers were reported, the detailed structure of the S2 C-terminus, particularly for those after amino acid (aa) 1146, was not resolved. We tested eight constructs of furin site mutated spike ectodomain with progressively truncated C-terminus in Chinese hamster ovary (CHO) cells. Interestingly, the C-terminal truncation to various positions between aa1152 and aa1192 resulted in higher expression levels and purification yields than the construct encompassing the entire ectodomain (S1208) (**Fig. 1A**). In addition, the C-terminal truncated spike proteins presented comparable, even better ACE2 binding activity to the trimeric StriFK protein (*17*) (**fig. S1A**). To minimize the potential epitope loss associated with C-terminal truncation, we finally chose the construct of S1192 encompassing aa 1-1192 (hereafter designated STFK) as an immunogen candidate for further study. As expected, the STFK was presented in monomeric form, as evidenced by the SEC-HPLC and native-PAGE analyses (**Fig. 1B**). In contrast to the trimeric StriFK, the STFK elicited significantly higher nAb titers against the pseudotyped virus (PsV) in mice at weeks 1 (*P* = 0.002) and 2 (*P* =0.028) after the 1^st^ prime vaccine dose, suggesting the advantage of the STFK for induction of rapid humoral response. After the 2^nd^ dose immunization, both STFK and StriFK-based vaccines generated comparable nAb titers (**Fig. 1C**).

**Fig. 1.**
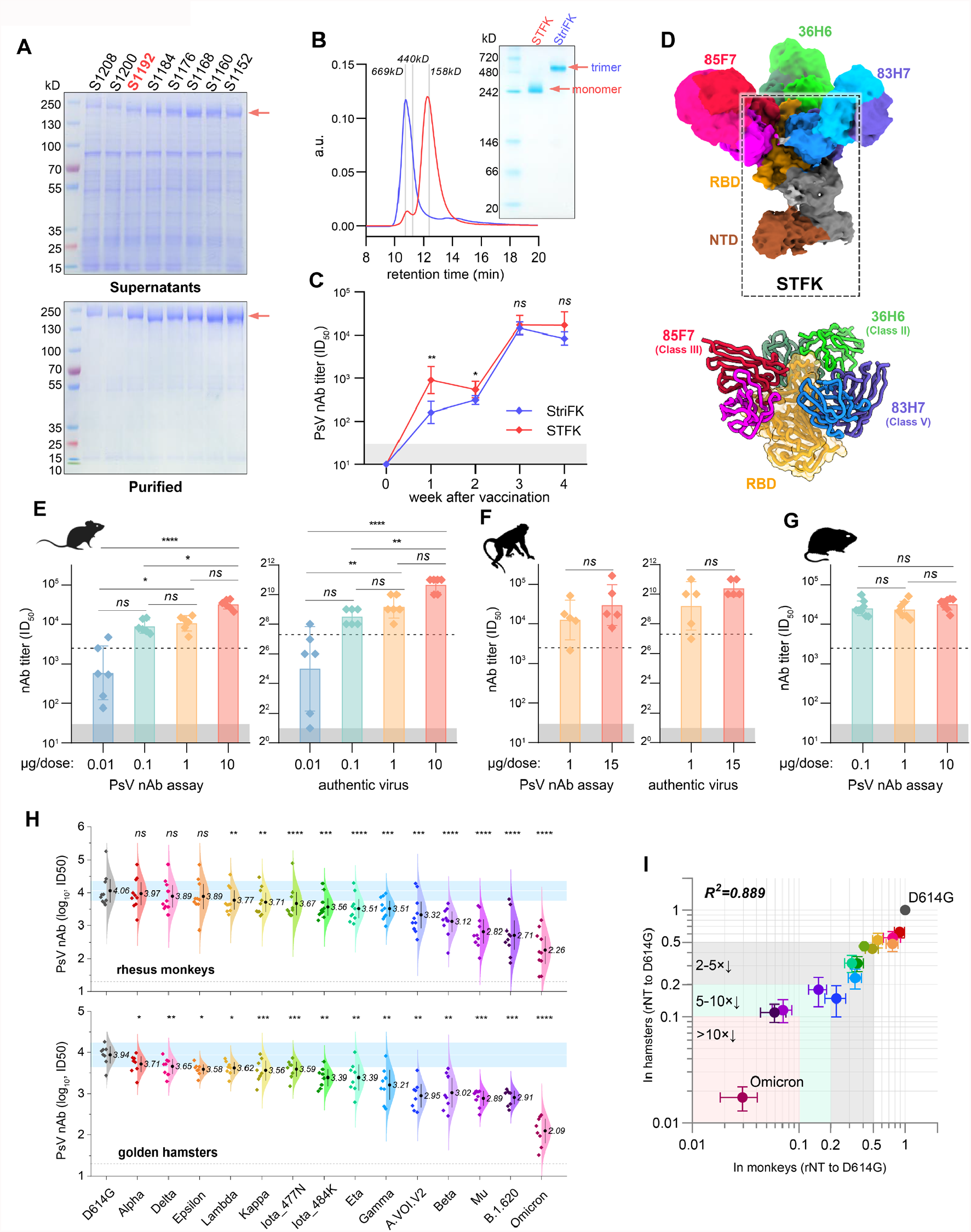
The monomeric STFK is highly immunogenic in rodents and nonhuman primates. **(A)** Reduced SDS-PAGE analyses for supernatants (top panel) and purified proteins (bottom panel) produced from constructs encoding progressive truncations from the C terminus of the furin site with mutated spike ectodomain in CHO cells. S1208, aa 1-1208; S1200, aa 1-1200; S1192, aa 1-1192; S1184, aa 1-1184; S1176, aa 1-1176; S1168, aa 1-1168; S1160, aa 1-1160; S1152, aa 1-1152. **(B)** Analyses of the monomeric STFK (aa 1-1192) and trimeric StirFK by SEC-HPLC (left panel) and Native-PAGE (right panel). **(C)** Comparison of the PsV nAb titers elicited by the STFK and StriFK in mice. BALB/c mice were immunized twice with 1 μg antigen at weeks 0 and 2. **(D)** 3.81 Å cryo-EM density map and corresponding atomic model of the STFK in complex of nAbs 36H6, 83H7, and 85F7. The black dotted box highlights the monomeric STFK protein. **(E-G)** Serum nAb titers against pseudotyped (left panel) and authentic SARS-CoV-2 viruses (right panel) of **(E)** BALB/c mice (n=6) or **(F)** Rhesus monkeys (n=5) received 2 shots of STFK vaccinations at different antigen doses. **(G)** Serum nAb titers against VSV-based PsV of hamsters vaccinated at 0.1 (n=8), 1 (n=7), or 10 μg (n=8) of STFK per dose. The immunization schedule was week 0/3 for **(E, G)** and week 0/4 for **(F)**. Sera were analyzed at weeks 4 **(E)**, 6 **(F)**, and 5 **(G)**. The dotted lines show the PsV nAb titers of WHO International Standard for anti-SARS-CoV-2 immunoglobulin using the same assays (NIBSC 20/136). **(H)** The nAb titers of sera from STFK-vaccinated rhesus monkeys (pooled of 1 and 15 μg groups, top panel) and hamsters (10 μg group, bottom panel) against lentiviral-pseudotyped SARS-CoV-2 spike variants compared to that against the ancestral D614G strain. The numbers showed the nAb GMT (log_10_) values. **(I)** Comparison of the cross-neutralizing activities of vaccinated hamsters (X-axis) and rhesus monkeys (Y-axis) against various lentiviral-pseudotyped SARS-CoV-2 variants. The relative nAb titer (rNT) was calculated as its ID_50_ ratio against a variant to the D614G control for each sample. Data in (D-H) were plotted as the geometric mean with SD. Dark shadows in (D-G) indicate the limit of detection (LOD). The dotted line in (H) indicates the LOD. Blue shadows in (H) represent the range of 50%-200% (within 2-fold changes) of the nAb GMT against D614G (as white line indicated). Uncorrected Kruskal-Wallis test (D, E, and G), Mann-Whitney *U* test (F), or Dunnett’s Multiple Comparison test (H) were used for intergroup statistical comparisons. Asterisks indicate statistical significance (\*\*\*\**P* < 0.0001; \*\*\**P* < 0.001; \*\**P* < 0.01; \**P* < 0.05; ns, not significant). Silhouettes indicating the species in (E-G) were from PhyloPic.org and available under the Public Domain Dedication 1.0 license.

To determine the structural basis for the excellent immunogenicity of the STFK, we resolved the cryo-electron microscopy (cryo-EM) structure of the STFK in complex with three previously reported nAbs 36H6, 83H7, and 85F7 (*18, 19*). Following the previous classifications of the nAbs targeting epitopes (Class I-V) (*20, 21*), the 36H6, 83H7, and 85F7 were categorized into Class II, V, and III, according to their binding modes, respectively (**fig. S2** and **S3**). The 3.81Å resolution structure of the immune-complex confirmed the monomeric form of the STFK, which could interact with three antigen-binding fragments (Fabs) of nAbs simultaneously (**Fig. 1D, fig. S2A**, and **Table S1**). The STFK is structurally similar to the monomeric form dissociated from a spike trimer (*22*). Due to the conformational flexibility in the monomeric form, the S2 subunit was not visualized in the reconstruction. However, in contrast to the trimeric spike, the STFK presents a more exposed RBD and NTD, thereby making the nAb epitopes more accessible and may contribute to its advantage for eliciting rapid nAb response.

Next, we evaluated the dose-dependent immunogenicity of the STFK-based vaccine with the FH002C adjuvant in the BALB/c mice, rhesus monkeys, and golden hamsters. In our previous study, the FH002C, a risedronate-modified new adjuvant, showed potent immunostimulatory effects for hormonal and cellular immune responses (*17*). In BALB/c mice, STFK vaccinations generated a dose-dependent response for the anti-spike IgG, anti-RBD IgG (**fig. S1B**), and neutralizing antibodies at 0.01 to 10 μg dose levels (**Fig. 1E**). Two injections of STFK at a dose level as low as 0.1 μg induced a potent nAb response showing geometric mean titers (GMT) of 3.9 log_10_ against the PsV and 362 against the authentic virus (200 TCID_50_) that were 3.5- and 2.3-fold higher than that of the NIBSC 20/136 anti-SARS-CoV-2 standard (1,000 IU/mL) in the corresponding assays (**Fig. 1E**). In rhesus monkeys, STFK vaccinations at 1 or 15 μg dose levels also elicited strong humoral immune responses (**Fig. 1F** and **fig. S1C**), as shown that immunized animals presented high nAb titers against either PsV (GMT=4.1 and 4.5 log_10_ for 1 and 15 μg groups, respectively) or authentic virus (GMT=588 and 1,351 for 1 and 15 μg groups, respectively). In addition, hamsters that received STFK vaccines of 0.1-10 μg per dose showed comparable and >4.0 log_10_ of nAb titers at week-2 after the boost dose (**Fig. 1G** and **fig. S1D**). Overall, the nAb titers (referred to as PsV nAb GMTs) elicited by 1 μg of FH002C-adjuvanted STFK vaccine were about 4.2 to 9.3-fold higher than that of the NIBSC 20/136 standard in mice, monkeys, and hamsters. Apart from humoral immunity, vaccinated mice also presented potent spike-specific T cell responses (*P <*0.001) (**fig. S1E**). These data demonstrated the promising immunogenicity of the monomeric STFK recombinant protein in rodents and nonhuman primates.

### Engineered STFK variant protein provides broad antigenic coverage which compensates with prototypic spike

We investigated the impacts of 14 VOC/VOI variants on nAbs raised by the prototypic STFK in animals. Notably, all sera from ten monkeys and eight hamsters at week-2 after 2-dose vaccinations showed detectable nAbs against all PsVs bearing VOC/VOI spike variants, including the newly emerged Omicron (**Fig. 1H**). However, by contrast to that against D614G lineage, STFK-elicited nAb titers in monkeys were markedly decreased (>5-fold) for Beta (6.5×), Mu (B.1.621) (14×), B.1.620 (17×), Omicron (34×); were mild to moderately reduced (2- to 5-fold) for A.VOI.V2 (4.4×), Gamma (2.9×), Eta (B.1.525) (3.2×), Iota_484K (B.1.526) (2.8×), Iota_477N (B.1.526) (2.4×), and Kappa (B.1.617.1) (2.0×) (**Fig. 1H** and **1I**). For Alpha, Delta, Epsilon (B.1.429), and Lambda (C.37) variants, the nAb titers only slightly changed a (<2-fold). Moreover, sera from immunized hamsters presented highly similar (R^2^=0.889, *P* < 0.001) cross-neutralizing profiles to that of monkeys (**Fig. 1I**). These results are consistent with findings in humans that the E484K-harboring variants and the Omicron may markedly evade nAbs raised by the prototypic spike.

Following the approach as graphically depicted in **Fig. 2A**, we aimed to develop a new STFK antigen providing complementary antigenic coverage to the prototypic protein to address the concerns for the evasive variants (**Fig. 2A**). As the Mu and Omicron variants had not emerged when our experiment started, we firstly tested mutated STFK antigens based on the spikes of Beta (STFK1351), Gamma (STFK1128), and B.1.620 (STFK1620) variants (**fig. S4**). Compared to those immunized with STFK, hamsters vaccinated with STFK1351, STFK1128, and STFK1620 showed 1.0-3.0×, 1.3-6.2×, and 1.7-5.5× increased nAb titers (GMTs) in neutralizing four immune-escape variants (Gamma, A.VOI.V2, Beta, and B.1.620) (**fig. S5**). The STFK1128 exhibited better immunogenicity than the other two, as it raised ∼4.0 log_10_ of nAb GMT in neutralizing its parental virus (Gamma) (**fig. S5**). In contrast, Beta variant-derived STFK1351 was poorly immunogenic.

**Fig. 2.**
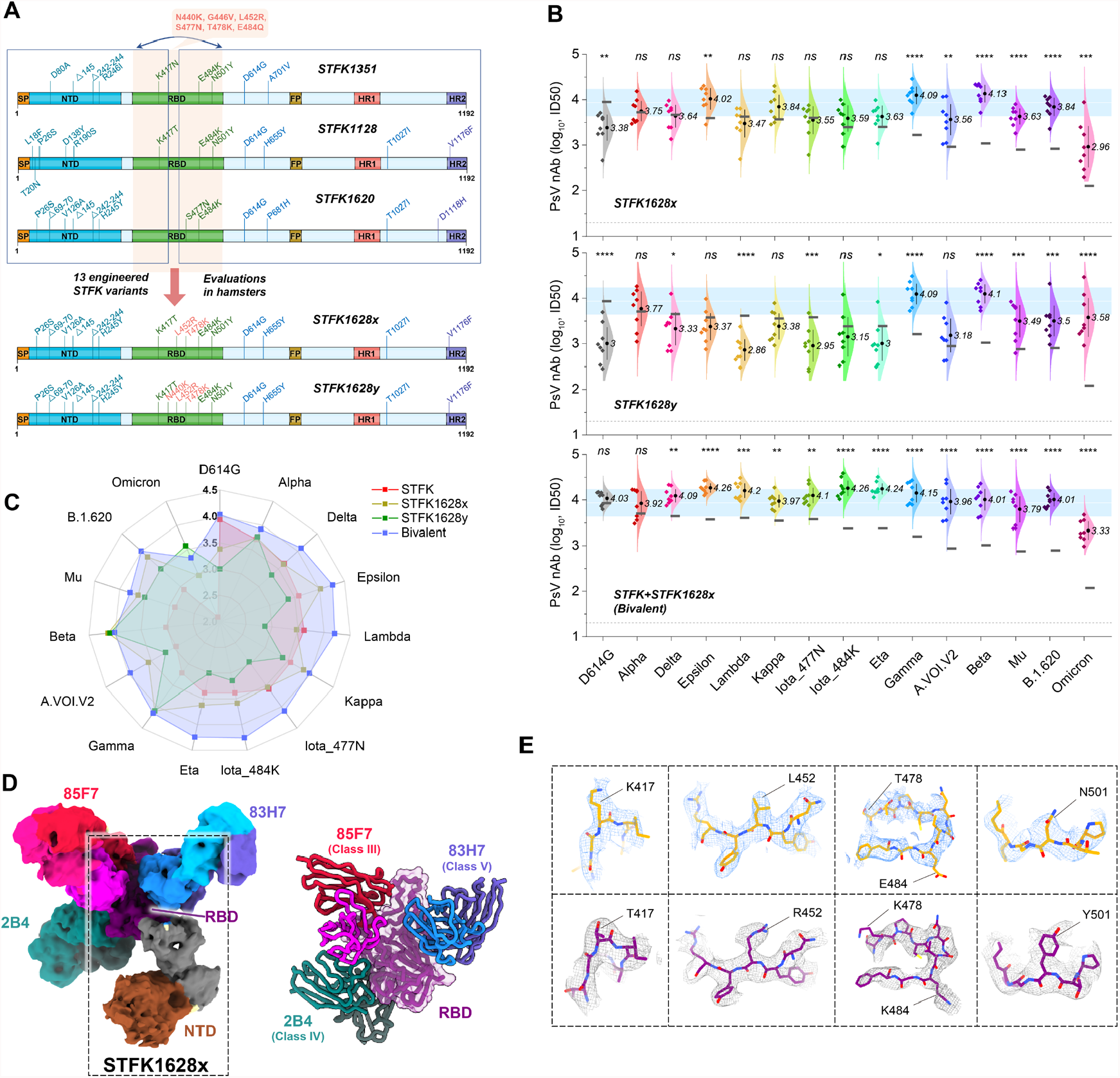
Neutralizing antibody responses elicited by engineered STFK variants in hamsters. **(A)** Schematic of the progressive approach to enlarge the cross-variants antigenic covering of a recombinant spike protein via inter-lineage chimera and mutation patch strategies. **(B)** The nAb titers of sera from hamsters (n=8) receiving vaccination of STFK1628x, STFK1628y, or a bivalent version of STFK+STFK1628x to neutralize lentiviral-pseudotyped SARS-CoV-2 variants. Dark horizontal lines indicate the nAb GMTs induced by the prototypic STFK vaccine against the corresponding variants. Blue shadows represent the range of 50%-200% (within 2-fold changes) of the nAb GMT against D614G (as white line indicated) induced by prototypic STFK. Dotted lines indicate the LOD (ID_50_=20) of the assay. Data were plotted as the geometric mean with SD. Uncorrected Fisher’s LSD tests were used for statistical comparisons (STFK v.s. modified STFK vaccine). Asterisks indicate statistical significance (\*\*\*\**P* < 0.0001; \*\*\**P* < 0.001; \*\**P* < 0.01; \**P* < 0.05; ns, not significant). **(C)** A spider plot showed the nAb GMTs (log_10_) against different SARS-CoV-2 variants of hamsters immunized with STFK, STFK1628x, STFK1628y, or the bivalent vaccine. Data were summarized from panel (B). **(D)** 3.88 Å cryo-EM density map and corresponding atomic model of the STFK1628x in complex of nAbs 83H7, 85F7, and 2B4. The black dotted box highlights the monomeric STFK1628x protein. **(E)** Comparisons of representative density maps of residues involved in RBD mutations on the STKF (top panel) and STFK1628x proteins (bottom panel).

We then introduced circulating RBD mutations absent in the Gamma variant but present in other VOC/VOI viruses into the STFK1128 backbone for the following engineering. We used the mutations of L452R (noted in Delta, Kappa, Epsilon), S477N (presented in Iota_477N and B.1.620), T478K (Delta-derived), and E484Q (Kappa-derived, to replace the E484K in STFK1128) to generate six new antigens (**fig. S6**). Hamster immunization tests revealed that the STFK1128e (L452R/S477N/E484Q), STFK1128f (L452R/T478K/E484K), and STFK1128g (L452R/T478K/E484Q) displayed improvements on the cross-neutralization spectrum in comparison to the STFK1128 (**fig. S6C**). As the Delta became the dominant variant worldwide since June 2021, we selected the STFK1128f exhibiting higher titers of nAbs to neutralize both Beta and Delta viruses as a candidate for further optimization.

Besides RBD mutations, NTD deletions presented in several VOC/VOI variants may also contribute to their immune-escape potentials. To cover the mutated NTD epitopes, we designed two inter-lineage chimeric constructs of STFK1328x and STFK1628x; the former included the NTD of STFK1351 (Beta), and the latter had the NTD of STFK1620 (B.1.620). Both constructs shared the RBD-S2 domain of STFK1128f, except a K417N in STFK1328x. Remarkably, the STFK1628x could elicit a broad and potent neutralizing antibody response in hamsters (**fig. S7A**), which showed 4.1-12.8× increased nAb titers (GMT) than prototypic STFK in neutralizing the A.VOI.V2 (4.1×), Gamma (7.6×), Beta (12.8×) and B.1.620 (8.5×). In contrast to its parental STFK1128f, the STFK1628x also exhibited higher nAb titers against most VOC/VOI variants (**Fig. 2B** and **fig. S7B**). These data supported the STFK1628x as a promising antigen candidate for the updated COVID-19 vaccine. As our previous study suggested that the aa 439-448 was another hot-spot region in addition to aa484 (*19*), we further made two modified STFK1628x versions, designated STFK1628y and STFK1628z, that included N440K and G446V, respectively (**Fig. 2B** and **fig. S7B**). The STFK1628y and STFK1628z displayed distinct antigenic profiles in hamsters (**fig. S7**). Notably, in contrast to the prototypic antigen, the STFK1628x and STFK1628y elicited significantly increased nAb in neutralizing two newly emerged variants, Mu (5.5×, *P* = 0.002 for STFK1628x; 4.0×, *P* = 0.010 for STFK1628y) and Omicron (7.4×, *P* = 0.002 for STFK1628x; 30.7×, *P* = 0.010 for STFK1628y) (**fig. S7C**). Following these data, we formulated a bivalent vaccine using the STFK1628x and the prototypic STFK at a mass ratio of 1:1. To most of the VOC/VOI variants, hamsters immunized with the bivalent vaccine showed significantly (*P* < 0.05) increased nAb levels to that elicited by the prototypic antigen in neutralizing the D614G virus (∼4.0 log_10_) (**Fig. 2B** and **2C**). Strikingly, the bivalent vaccine yielded a nAb GMT of 2,130 (ID_50_ range: 9,61 to 4,763) to the highly immune-evasive Omicron, which was about 36-fold higher than the NIBSC 20/136 immunoglobulin standard (ID_50_=60 to Omicron). Taken together, STFK plus STFK1628x provided a full-spectrum neutralization coverage to all VOC/VOI variants.

We also obtained a 3.88 Å cryo-EM structure of the STFK1628x in complexed with three nAbs 83H7, 85F7, and 2B4 (**Fig. 2D, fig. S2B**, and **Table S1**). As the T478K abolishes the activity of 36H6 nAb, we replaced the 36H6 with a class IV mAb of 2B4 with cross-SARS-CoV-1/2 neutralization potency (**fig. S3D**) (*18*). As expected, the STKF1628x presented a similar structure to STFK but showed distinguished densities on the mutation sites, such as 417, 452, 478, 484, and 501, corresponding to its alternative antigenic profile (**Fig. 2E**).

### The bivalent vaccine protects hamsters against intranasal SARS-CoV-2 challenges

To assess the ability of the STFK-based vaccine to mediate protection against SARS-CoV-2, we intranasally challenged hamsters that received STFK, STFK1628x, or bivalent vaccines (**Fig. 3A**). For either challenge with the ancestral strain or Beta variant, vaccinated hamsters showed an average of 2.2-4.6% weight increase to their baseline levels by the end of a 7-day follow-up (**Fig. 3B** and **3C**). By contrast, unvaccinated animals showed a maximum weight loss of 14.8% and 13.8% by 7 days post-infection (dpi) in the ancestral strain and Beta variant challenges, respectively. Moreover, 1 of 8 and 5 of 8 animals died from ancestral SARS-CoV-2 and beta variant infections, respectively, but none in the vaccinated groups (**Fig. 3D** and **3E**).

**Fig. 3.**
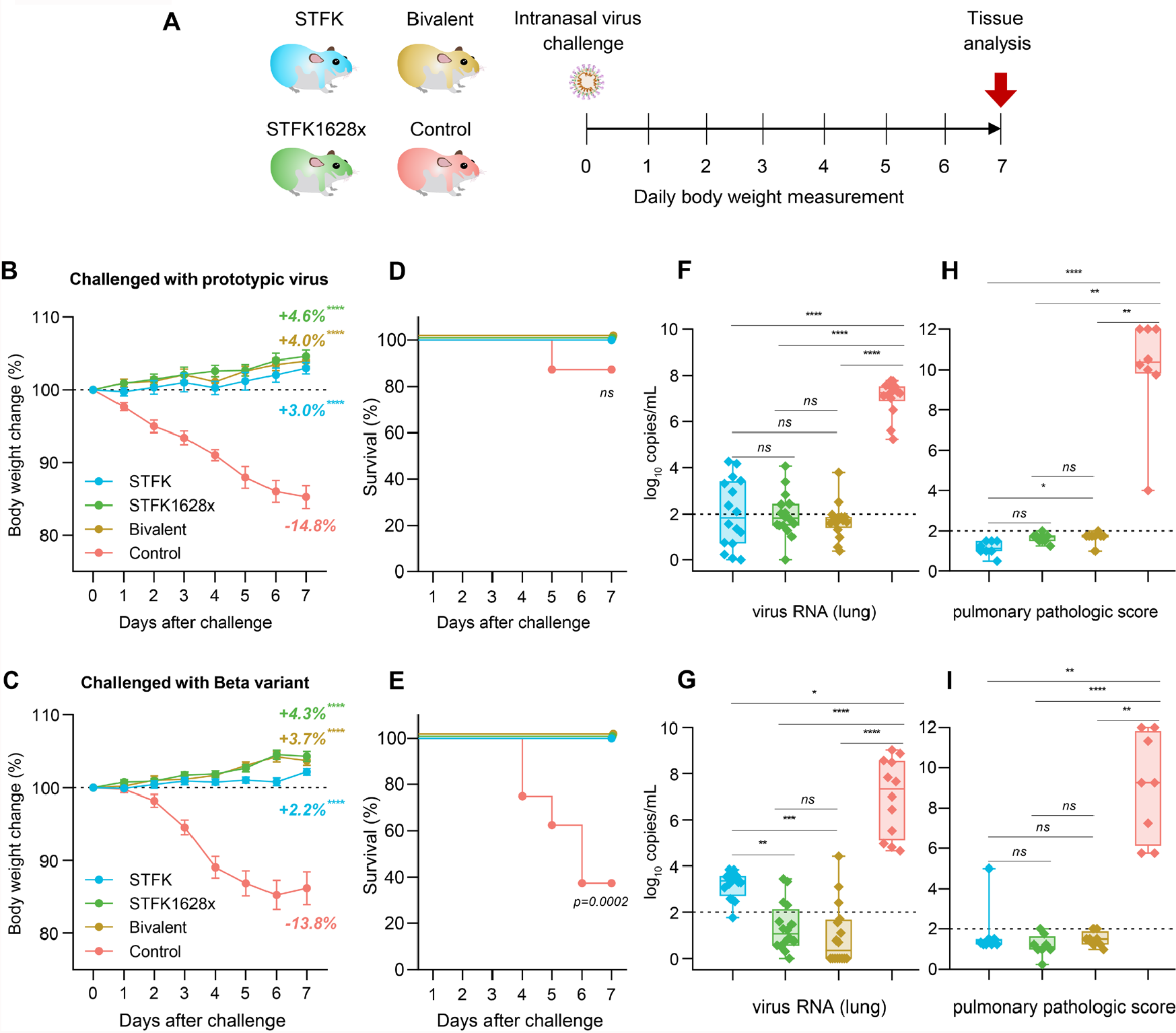
The STFK, STFK1628x, and bivalent vaccines offer protection against ancestral SARS-CoV-2 and Beta variant intranasally challenged in hamsters. **(A)** Schematic representation of the intranasal virus challenge evaluation for vaccine effectiveness. A total of 64 hamsters were used for two independent tests. Each group (indicated in different colors) included eight hamsters (4 males and 4 females) that received 2-dose vaccinations (at weeks 0/3) before virus challenges. Hamsters were intranasally challenged with 1×10^4^ PFU of ancestral SARS-CoV-2 or Beta variant (B.1.351) at week 5. After a 7-day weight monitoring follow-up, animals were euthanized for tissue analyses. Weight changes (**B, C**), survival curves (**D, E**), lung viral RNA levels (in two independent lung tissues, LOD=2 log_10_ copies/mL) (**F, G**), and pulmonary pathological scores (**H, I**) of hamsters challenged by ancestral SARS-CoV-2 **(B, D, F, H)** or beta variant **(C, E, G, I)** were shown. Data in (**B, C**) were plotted as means±SEM. Data in (**F-I**) were shown as box and whisker plots; the median, first quartile, third quartile, minimum, and maximum values were plotted. Dunnett’s Multiple Comparison test (**B, C)**, two-sided log-rank test (**D, E**), or uncorrected Kruskal-Wallis test (**F-I**) were used for intergroup statistical comparisons. Asterisks indicate statistical significance (\*\*\*\**P* < 0.0001; \*\*\**P* < 0.001; \*\**P* < 0.01; \**P* < 0.05; ns, not significant).

At 7 dpi, the median viral RNA levels of control hamsters challenged by the prototypic virus were 7.26 (range 5.24-7.78) log_10_ in the lung, 6.88 (range 6.12-7.32) log_10_ in the nasal turbinate, and 6.29 (range 5.04-6.67) log_10_ copies/mL in the trachea (**Fig. 3F** and **fig. S8A**). By contrast, hamsters that received vaccinations of either STFK, STFK1628x, or the bivalent version showed significant (*P* < 0.01 for each comparison) viral RNA reductions by >5.0 log_10_, 2.0-3.0 log_10,_ 3.0-4.0 log_10_ copies/mL in tissues of the lung, nasal turbinate, and trachea, respectively (**Fig. 3F** and **fig. S8A**). To protect the prototypic virus challenge, the three vaccine candidates appeared with comparable efficacy (*P* > 0.05) in decreasing viral loads of the respiratory tract tissues (**Fig. 3F** and **fig. S6A**). In the Beta variant challenges, control hamsters also showed high levels of viral RNA similar to that infected with the prototypic virus in their respiratory tract tissues. For vaccinated animals, the medians of pulmonary viral loads were 1.05, 0.35, and 3.36 log_10_ copies/mL in the STFK1628x, bivalent and STFK groups, corresponding to reductions of 6.27 (*P* < 0.0001), 6.96 (*P* < 0.0001), and 3.96 (*P* = 0.01) log_10_ to controls, respectively (**Fig. 3G**). In tissues of nasal turbinate and trachea, no statistically significant difference in viral RNA suppression was observed among the 3 vaccination groups (**fig. S8B**), but hamsters immunized with the bivalent vaccine presented relatively lower viral loads than the others.

In addition to mediating virological suppressions, the three vaccines also protected hamsters from lung disease caused by SARS-CoV-2 infections. Pathological examinations revealed most unvaccinated hamsters presented severe pulmonary diseases at 7 dpi regardless of the challenged virus type (**Fig. 3H** and **3I, fig. S8C** and **S8D**). In contrast, gross lung observations and pulmonary pathology scorings demonstrated all vaccinated animals were free from moderate-to-severe pneumonia, with the only exception noted in one from the STFK group challenged with Beta variant (**Fig. 3I**). These data indicated that the STFK, STFK1628x, and the bivalent vaccines effectively protected hamsters against the SARS-CoV-2 challenge. Moreover, the STFK1628x and the bivalent vaccine showed better protective efficacy against the Beta variant challenge in hamsters.

### The bivalent vaccine blocks SARS-CoV-2 transmissions in hamsters

In addition to protecting vaccine users from SARS-CoV-2 infection and pathology, an ideal COVID-19 vaccine should also reduce viral shedding and transmission by vaccinated individuals exposed to the virus. For assessments, index hamsters were immunized with either STFK1628x or the bivalent vaccine and were challenged intranasally with ancestral SARS-CoV-2 or Beta variant. One day later, naïve hamsters as sentinels were cohoused with index animals for 24 hours **(Fig. 4A**). After a subsequent 7-day follow-up, sentinels of the unvaccinated index hamsters showed an average weight loss of 4.0% and 1.7% in the ancestral virus and beta variant challenges, respectively. In contrast, sentinels of vaccinated indexes receiving either the STFK1628x or the bivalent version exhibited gradually increased weights (**Fig. 4B** and **4C**). By the end of the experiment, all sentinels of the unvaccinated indexes had detectable viral RNA with approximate levels of 6.0-7.0 log_10_ copies/mL in the respiratory tract tissues (**Fig. 4D** and **4E**). By contrast, 4 (50%) and 1 (12.5%) sentinels cohoused with STFK1628-vaccinated hamsters, when challenged with the Beta variant and the ancestral SARS-CoV-2 respectively, showed detectable viral RNA in either tissue from the lung, nasal turbinate, or trachea. Remarkably, for either challenge of the two viruses, no sentinel hamster of the indexes immunized with the bivalent vaccine showed detectable viral RNA in any tissues from the lung, nasal turbinate, and trachea (**Fig. 4D** and **4E**). These data supported complete protection from SARS-CoV-2 transmission conferred by the bivalent vaccine in hamsters.

**Fig. 4.**
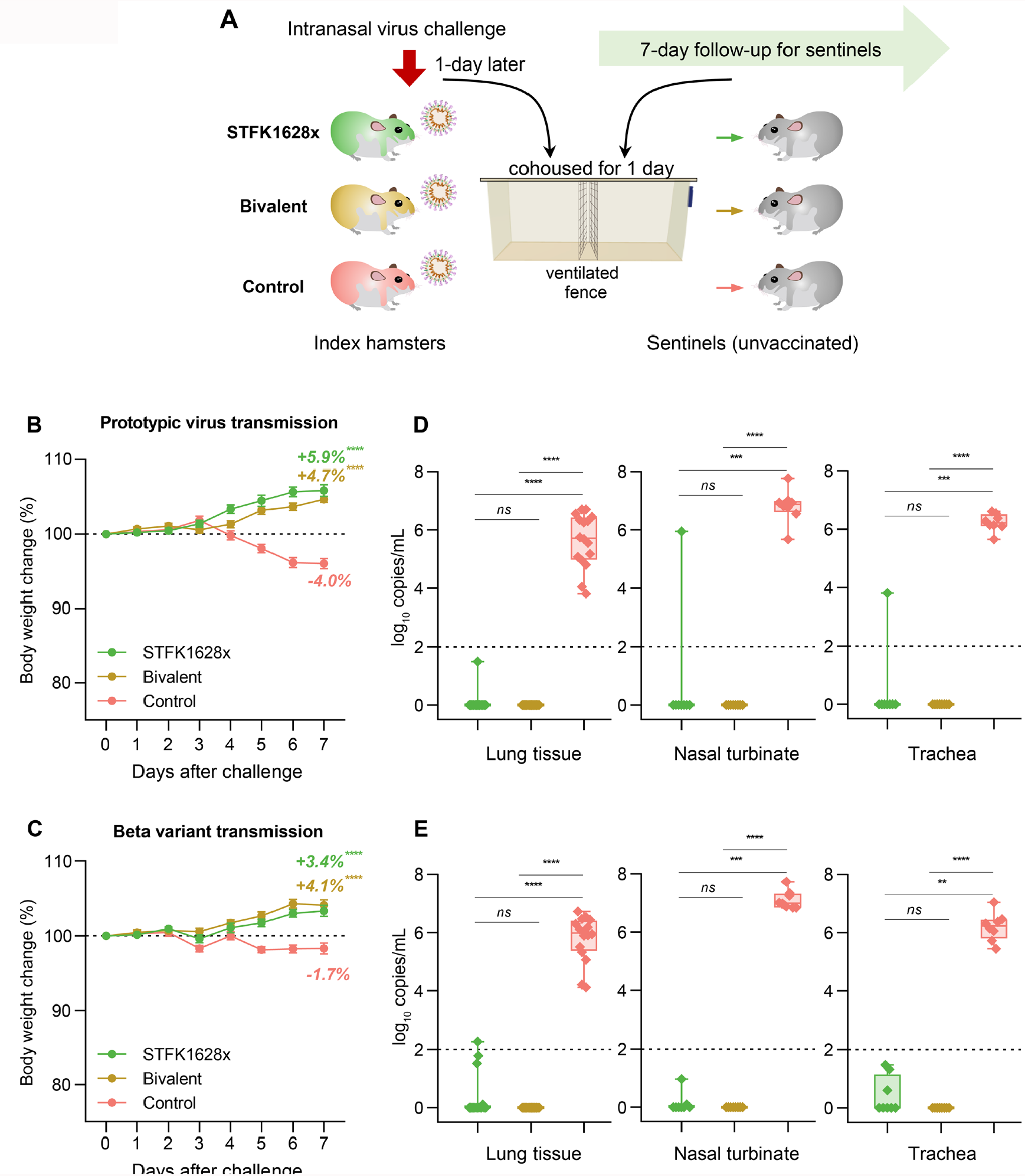
The monovalent STFK1628x and the STFK/STFK1628x-combined bivalent vaccines prevent SARS-CoV-2 transmission among hamsters. **(A)** Schematic diagram of the experimental design. Vaccinated-(STFK1628x or bivalent vaccine) and unvaccinated-index hamsters (n=4) were intranasally challenged with 1×10^4^ PFU of ancestral SARS-CoV-2 or Beta variant. One day later, each 2 index hamsters were cohoused with 4 naïve sentinels for one day (separated by a double-layer ventilated fence in the same cage). Sentinel hamsters (n=8) were followed for 7-day and then euthanized for tissue analyses. Weight changes **(B, C)** and viral RNA levels in the respiratory tract tissues **(D, E)** of sentinels after cohoused with index hamsters challenged by ancestral SARS-CoV-2 **(B, D)** or beta variant **(C, E)** were shown. Viral RNA levels in tissues of the lung (in two independent tissues), nasal turbinate, and trachea were measured by qRT-PCR (LOD=2 log_10_ copies/mL). Data in (B and D) were plotted as means ± SEM. Data in (C and E) were shown as box and whisker plots; median, first quartile, third quartile, minimum value, and maximum value were plotted. Dunnett’s Multiple Comparison test (**B, C)**, or uncorrected Kruskal-Wallis test (**D, E**) were used for intergroup statistical comparisons, and asterisks indicate statistical significance (\*\*\*\**P* < 0.0001; \*\*\**P* < 0.001; \*\**P* < 0.01; \**P* < 0.05; ns, not significant).

## Discussion

With the widespread SARS-CoV-2 infections in large populations of humans and other susceptible animals, the emergence of antigenic drift variants seems inevitable. The newly identified Omicron variant with >30 spike mutations brought great challenges to established immunity by vaccination or resolved infections. Recent studies revealed that the Omicron had led to widespread escape from nAbs acquired from infections by the ancestral SARS-CoV-2 or other VOC variants, including Alpha, Beta, Gamma, and Delta (*23, 24*), exhibiting a distinctly altered antigenicity. This variant is also highly resistant against nAbs elicited by double immunizations of the authorized COVID-19 vaccine (*25*). Though some studies have found the 3^rd^ dose of vaccination may increase the nAb titer against Omicron, it was still lower than that against other variants (e.g., Delta) by several folds (*26, 27*). Next-generation COVID-19 vaccines with pan-variants protection are urgently required to deal with the growing threat of Omicron-like immune-escape variants.

Adding another spike antigen of an immune-escape variant is the most common approach for developing an updated COVID-19 vaccine. Before Omicron emergence, the Beta and Gamma were two noticeable VOCs with increased immune evasion concerns; therefore, most previous studies used spikes from one of the two variants as the added antigen. The mRNA-1273.351, a Beta variant-based mRNA vaccine developed by Moderna, could elicit a potent nAb response against the Beta variant in mice but was inferior to that derived from the prototype-based mRNA-1273 in neutralizing the Gamma and Epsilon variants (*27*). Neutralizing antibodies elicited by vaccines formulated with either Beta or Gamma-derived spikes in animals appeared to be less effective against the Delta variant (*28, 29*). Consistent findings were also noted in the presented study (fig. S5). Moreover, in our data, neither spike antigens from variants of Beta (STFK1351), Gamma (STFK1128), nor B.1.620 (STFK1620) induced markedly improved nAb against Omicron in hamsters (fig. S7). Therefore, it is unlikely to generate broad-spectrum antigenic coverage by using a pre-existing naturally occurring variant.

Here, we provided a progressive approach to enlarge the cross-variants antigenic covering of a recombinant spike protein. This approach followed inter-lineage chimera and mutation patch strategies. As the NTD, RBD, and S2 contain neutralizing antibody-targeting epitopes, inter-lineage chimeric spike antigens might confer cross-variants antibody response (*30*). The engineered STFK1628x, providing mutually-complementary antigenic coverage to the prototypic spike, was a chimeric construct of the B.1.620-derived NTD and the Gamma variant derived RBD-S2. The B.1.620 NTD harbored amino-acid deletions of Δ69-70, Δ145, and Δ242-244, correspond to changed epitopes on the N2, N3, and N5 loops (*31*). Notably, Alpha, Eta, and Omicron variants shared the Δ69-70 and Δ145, whereas the Beta variant contained Δ242-244. The RBD-S2 of the two engineered antigens were derivates of the Gamma spike with additional antigenic mutation patches of L452R and T478K. Our and others’ studies both suggested the Gamma-derived spike was highly immunogenic to elicit potent nAb response against E484K-harboring variants, such as Beta and Gamma (*32*). As two Delta-derived characteristic mutations, the L452R and T478K introduction showed added value to improve the neutralization response against the Delta variant (fig. S6). In contrast to STFK, the monovalent STFK1628x and the STFK/STFK1628x-combined bivalent vaccines conferred improved protection against viral infection and transmission of SARS-CoV-2 Beta variant in hamsters (Fig. 3 and 4). Given the potent cross-variants neutralization response elicited by the STFK/STFK1628x-combined bivalent vaccine in hamsters, we can expect the broad-spectrum protection of the new vaccine.

Remarkably, the STFK1628x and its STFK1628y derivate elicited significantly higher nAb titers against the Omicron variant than the prototypic STFK (fig. S7D), possibly attributed to the shared mutations of Δ69-70, K417N/T, T478K, E484K/A, and N501Y. In comparison to STFK1628x, the STFK1628y has an additional N440K mutation but presented a 4-fold higher nAb titer to neutralize Omicron (fig. S7D), suggesting the N440K plays an essential role in changing the antigenicity of this variant. Notably, as both antigens were generated before Omicron emergence, these results were encouraging and suggested the possibilities of prospective antigen design and vaccine preparations against an unknown future variant. Timely and comprehensive assessments for the antigenic influence of newly emerged spike mutations are essential for guiding vaccine antigen design.

Although trimerized spike antigens were commonly used in current COVID-19 vaccines, our study found that the monomeric spike protein is also highly immunogenic in rodents and nonhuman primates. The CHO-derived C-terminal truncated STFK proteins maintained comparable ACE2 binding activity as the trimeric StriFK, conferred potent immune responses, and complete protection against SARS-CoV-2 in hamsters. The monomeric form with more exposed protein surfaces may enable more accessibilities of cryptic epitopes and thereby facilitating the rapid and diverse antibody response. Removing the additional motif required for protein trimerization eliminates unexpected immune responses targeting the trimeric domain. In addition, the high yield of STFK in the CHO-cell expression system (800-1,000 mg/L from stable cell lines) is beneficial to reduce manufacturing costs and deal with the global shortage of COVID-19 vaccines. In summary, our study provides a new way to design new antigens for next-generation COVID-19 vaccines aiming to confer broad-spectrum protection.

## Materials and Methods

### Constructs, protein expressions, and purifications

Trimeric S-ectodomain proteins of SARS-CoV-2 (StriFK) were expressed and purified as previously described (*17*). Expression cassettes encoding monomeric S-ectodomain proteins (containing mutated furin-site, RRAR to GSAS) involved in the study were generated by site-directed mutagenesis via PCR cloning based on the parental codon-optimized StriFK (*17*). The expression cassettes of S1152 to S1208 with C-terminal 8×His-tag were cloned into the EIRBsMie vector (19). The tag-free STFK and STFK variants were constructed into a pGS01b vector, modified from the pCGS3 vector containing glutamine synthetase (GS) selection marker (Sigma Aldrich). All STFK constructs had an N1192M modification to reduce potential protein aggregation, possibly attributed the K933-N1192 interaction (*33*). As previously described, transient protein expressions were performed using the ExpiCHO expression system (Thermo Fisher Scientific). Stable cell lines expressing the STFK, STFK1628x, and STKF1628y were generated via transfections of the PGS01b-vectored constructs into CHOZN GS^-/-^ cells (Sigma Aldrich) following GS selections and single-cell clonings. Polyhistidine-tagged proteins (S1152 to S1208) purified from culture supernatants were collected on day 7 after transfection using Ni Sepharose 6FF (Cytiva). The tag-free STFK proteins were purified by using Q-FF Sepharose ion-exchange chromatography (Cytiva). Recombinant human ACE2 (human Fc tag, rACE2) protein also was produced in ExpiCHO-S cells and purified by protein-A affinity chromatography column (Cytiva) as previously described (*17*).

### SDS-polyacrylamide gel electrophoresis (SDS-PAGE) and Native-PAGE

SDS-PAGE analyses were performed using 4-12% SurePAGE (Genscript), the precast mini polyacrylamide gels. For native-PAGE, the protein samples were mixed with the Native Sample Buffer (BIO-RAD) in equal volume and then subjected to electrophoresis using the 7.5% Mini-Protean TGX Precast Protein Gels (Bio-RAD) in a non-denaturing buffer. Gel images were captured using FUSION FX7 Spectra multispectral imaging system (Vilber).

### Size exclusion chromatography (SEC-HPLC)

The SEC-HPLC analysis shown in Fig. 1B was performed using a TSK-GEL G3000PWXL column on an HPLC system (Waters Alliance) and conducted as described previously(*17*).

### ACE2 binding assays

A capture antibody 45C3 (developed in our laboratory) recognizes spike S2 domain was coated in 96-well microplates at 200 ng per well. Plates were incubated overnight at 4 °C and then blocked with ELISA-blocking buffer (Wantai BioPharm). The STFK proteins were twofold serially diluted from 10 μg mL^−1^ to 9.8 ng mL^−1^ in duplicate and then added to wells (100 μL). After incubation for 1 hour at 25°C followed by washing with PBST buffer, rACE2 protein solution (100 μL per well, 1 μg mL^−1^) was added to the wells. Subsequently, the microplates were incubated for 1 hour at 25°C. After washing five times, HRP-conjugated anti-human IgG (Thermo Fisher Scientific) solutions were added and incubated for 1 hour at 25°C. Following washing five times, tetramethylbenzidine chromogen (TMB) solution (Wantai BioPharm) was added into microplates 100 μL per well. After a further 10 minutes of incubation at 25°C, 2 M H_2_SO_4_ was added to stop the chromogen reaction, and the optical density (OD_450-630_) value was measured. The half-maximal effective concentration (EC_50_) was calculated by the 4-parameter logistic (4PL) regression using GraphPad Prism 8 software.

### Vaccine preparations

Recombinant spike protein subunit vaccines used in this study were composed of spike proteins and a nitrogen bisphosphonate-modified zinc-aluminum hybrid adjuvant (FH002C), which was described detailly in our previous study(*17*). Briefly, the proteins were mixed with an equal volume of 2× concentration FH002C adjuvant to achieve the final desired concentration of antigen in the final formulation. All vaccine formulations were mixed well and stored at 2-8 °C until use.

### Experimental animals

BALB/c and C57BL/6 mice were purchased from Shanghai SLAC Laboratory Animal Co., Ltd. Lakeview Golden (LVG) Syrian hamsters were purchased from Charles River Laboratories (Beijing). The animals were fed in Specific-pathogen-free circumstances. The mouse and hamster studies were carried out in strict accordance with the recommendations of the Guide for the Care and Use of Laboratory Animals under the approval of the Institutional Animal Care and Use Committee of Xiamen University. The rhesus monkey experiment was conducted at the Key Laboratory of Technical Evaluation of Fertility Regulation for Nonhuman Primate Inc in Fujian province.

### Mouse immunizations

Six to eight-week-old BALB/c mice (n=6 per group) were immunized with STFK vaccines at 0.01, 0.1, 1, or 10 μg per dose in 150 μL through intramuscular injection following a two-dose schedule at weeks 0 and 3. Sera were collected at week 4 via retro-orbital bleeding to measure antibody titers. For T cell response assessment, six to eight-week-old C57BL/6 mice were immunized with STFK vaccine at 10 μg per dose in 150 μL through intramuscular injection. Immunized mice were sacrificed on day 7 after immunization to collect splenocytes for further assay.

### Rhesus monkey immunizations

Ten rhesus monkeys were allocated randomly into two groups (three females and two males per group). Groups of monkeys were injected with 1 μg or 15 ug of STFK vaccine per dose (150 μL) via the intramuscular route for 2 doses. All monkeys were vaccinated at weeks 0 and 4. Two weeks after boosting, serum samples were collected for antibody analyses, including measurement of anti-spike IgG, anti-RBD IgG, pseudovirus neutralizing antibody, and authentic neutralizing antibody titers.

### Hamster immunizations

Six to eight-week-old hamsters were used to evaluate the immunogenicity and protective effect of the vaccine candidates. Each group contained four males and four females. Groups of hamsters were immunized intramuscularly twice with the vaccine candidates at 10 μg per dose in 200 μl, three weeks apart. All serum samples were collected at week-2 after the 2^nd^ dose via retro-orbital bleeding to measure the antibody titers.

### Pseudovirus (PsV) neutralization assays

The nAb titers against the ancestral spike-pseudotyped virus presented in Fig. 1C and 1E-1G were determined by using a vesicular stomatitis virus (VSV) system as previously described (*34*). The International Standard for anti-SARS-CoV-2 immunoglobulin (NIBSC code: 20/136) was obtained from National Institute for Biological Standards and Control, UK (*35*).

The lentiviral-based pseudovirus (LV) neutralization assay was used to determine vaccine-elicited nAbs against circulating SARS-CoV-2 variants (Fig. 1H, 2, S5, S6, and S7). The lentiviral pseudoviruses bearing spikes from SARS-CoV-2 variants were generated as described previously (*19, 36*), including D614G (site-directed mutagenesis), Alpha (B.1.1.7, GISAID accession number: EPI_ISL_601443), Beta (B.1.351, EPI_ISL_700428), Gamma (P.1, EPI_ISL_792680), Delta (B.1.617.2, EPI_ISL_1662451), Omicron (B.1.1.529, BA.1, EPI_ISL_6704867), Iota_484K (B.1.526_484K, EPI_ISL_1009654), Iota_477N (B.1.526_477N, EPI_ISL_995145), Epsilon (B.1.429, EPI_ISL_873881), Eta (B.1.525, EPI_ISL_762449), Kappa (B.1.617.1, EPI_ISL_1595904), A.VOI.V2 (EPI_ISL_1347941), Lambda (C.37, EPI_ISL_2921532), Mu (B.1.621, EPI_ISL_3933281) and B.1.620 (EPI_ISL_1620228). The PsV nAb measurements were performed as previously described using Opera Phenix or Operetta CLS High-Content Analysis System (PerkinElmer) (*19*).

### Authentic SARS-CoV-2 neutralization assay

The nAb titers of sera from immunized animals against authentic SARS-CoV-2 were detected using a cytopathic effect (CPE)-based microneutralization assay as previously described (*37*). The ancestral virus (BetaCoV/Jiangsu/JS02/2020, EPI_ISL_411952) was used. Briefly, serum samples were twofold serially diluted from 1:4 to 1:8192 in duplicate with DMEM medium. All prepared samples were mixed with the virus of 200 TCID_50_ and incubated for 2 hours at 37 °C. The mixtures (150 μL per well) were added to a monolayer of Vero cells in a 96-well plate and incubated at 37 °C supplying with 5% CO2. Three-day later, the cytopathic effect was assessed with microscopic examinations. The neutralizing titer of serum was expressed as the reciprocal of the maximal sample dilution that protects at least 50% of cells from CPE.

### Enzyme-linked immunospot (ELISpot) assay

According to the manufacturer’s instructions, the assays were performed with mouse IFN-γ ELISpot plates kits (Dakewe Biotech, 2210005). In brief, single-cell suspensions were obtained from mouse spleen (10^6^ cells per well) through grinding in 70 μm cell strainers and were seeded in anti-mouse IFN-γ antibody precoated ELISpot plates. Then, cells were incubated with pooled peptides of SARS-CoV-2 spike (15-mer peptides with 11aa overlap covering the entire spike protein; GenScript) and cultured at 37°C with 5% CO_2_ for 20 hours. Spots were counted and analyzed by using CTL-ImmunoSpot S5 (Cellular Technology Limited). The numbers of IFN-γ-secreting cells were calculated by subtracting phosphate-buffered saline (PBS)-stimulated wells from spike peptide pool-stimulated wells.

### Anti-RBD, anti-spike IgG measurements

Microplates pre-coated with recombinant antigens of RBD or spike ectodomain were provided by Beijing Wantai Biological Pharmacy. The measurements were performed following previously described procedures (*17*), with the only difference that the cutoff (CO) value was set as 0.1 (OD_450-630_). The IgG titer of each sample was determined as the cutoff index (OD_450-630_/CO) at the dilution limit multiplied by the maximum dilution folds. Representative data from technical replicates were performed at least twice for plotting.

### Cryo-EM sample preparation and data collection

Fabs of 36H6, 83H7, 2B4, and 85F7 were prepared by papain digestion of the mAbs and further purified with MabSelect SuRe (Cytiva). Aliquots (3 μl) of 3.5 mg/mL mixtures of purified STFK or STFK1628x proteins in complex with excess Fab fragments of nAbs were incubated in 0.01% (v/v) Digitonin (Sigma) and then loaded onto glow-discharged (60 s at 20 mA) holey carbon Quantifoil grids (R1.2/1.3, 200 mesh, Quantifoil Micro Tools) using a Vitrobot Mark IV (ThermoFisher Scientific) at 100% humidity and 4°C. Data were acquired using the SerialEM software on an FEI Tecnai F30 transmission electron microscope (ThermoFisher Scientific) operated at 300 kV and equipped with a Gatan K3 direct detector. Images were recorded in the 36-frame movie mode at a nominal 39,000× magnification at super-resolution mode with a pixel size of 0.339 Å. The total electron dose was set to 60 e^−^ Å^−2^, and the exposure time was 4.5 s.

### Image processing and 3D reconstruction

Drift and beam-induced motion correction were performed with MotionCor2 (*38*) to produce a micrograph from each movie. Contrast transfer function (CTF) fitting and phase-shift estimation were conducted with Gctf (*39*). Micrographs with astigmatism, obvious drift, or contamination were discarded before reconstruction. The following reconstruction procedures were performed by using Cryosparc V3 (*40*). In brief, particles were automatically picked by using the “Blob picker” or “Template picker”. Several rounds of reference-free 2D classifications were performed, and the selected good particles were then subjected to ab-initio reconstruction, heterogeneous refinement and final non-uniform refinement. The resolution of all density maps was determined by the gold-standard Fourier shell correlation curve, with a cutoff of 0.143. Local map resolution was estimated with ResMap (*41*).

### Atomic model building, refinement, and 3D visualization

The initial models of nAbs were generated from homology modeling by Accelrys Discovery Studio software (available from: https://www.3dsbiovia.com). The structure from the prototypic trimeric spike (PDB no. 6VSB) (*42*) was used as the initial modes of our proteins. We initially fitted the templates into the corresponding final cryo-EM maps using Chimera (*43*), and further corrected and adjusted them manually by real-space refinement in Coot (*44*). The resulting models were then refined with phenix.real_space_refine in PHENIX (*45*). These operations were executed iteratively until the problematic regions, Ramachandran outliers, and poor rotamers were either eliminated or moved to favored regions. The final atomic models were validated with Molprobity (*46, 47*). All figures were generated with Chimera or ChimeraX (*48*).

### SARS-CoV-2 virus challenges in hamster

Two weeks after boosting, hamsters were challenged with the ancestral SARS-CoV-2 of hCoV-19/China/AP8/2020 (EPI_ISL_1655937) or Beta variant of hCoV-19/China/AP100/2021 (EPI_ISL_2779639). For the intranasal challenge, hamsters were challenged with 1×10^4^ PFU of SARS-CoV-2 virus (diluted in 100 μL of PBS) through the intranasal route under anesthesia. For virus transmission-blocking study, vaccinated- or unvaccinated-hamsters were intranasally inoculated with 1×10^4^ PFU of ancestral SARS-CoV-2 or Beta variant as indexes. One day post-infection, index hamsters were cohoused with naïve sentinels for one day (Fig. 4A). The daily diet was limited to 7 g per 100 g of body weight to prevent animals from overeating. All hamsters were monitored for body weight until being humanely euthanized on day 7 after challenge or exposure. The respiratory tissues of hamsters were collected for viral RNA quantification or histopathological assessments. All challenge experiments were conducted in the Animal Biosafety Level 3 (ABSL-3) facility.

### SARS-CoV-2 RNA quantification

Viral RNA levels in tissues from hamsters were determined by SARS-CoV-2 RT-PCR Kit (Wantai BioPharm). For each animal, two pieces (separately) of lung tissue (0.1∼0.2 g each), one piece of the trachea tissue (0.1∼0.2 g), and a half of nasal turbinate (0.8∼1.2 g) were respectively homogenized with TissueLyser II (Qiagen) in 1 ml PBS. Viral RNA in tissue lysates was extracted using the QIAamp Viral RNA Mini Kit (Qiagen) and subjected to qRT-PCR assays. Representative data from technical replicates were obtained from at least two independent experiments for plotting.

### Histopathology

The lung tissues from challenged hamsters were fixed with neutral buffered formalin for 48 hours and processed routinely into paraffin blocks. Then tissues were sectioned to 3 μm by microtome (Leica). Next, the fixed lung sections were stained with hematoxylin and eosin (Maxim Biotechnology). Whole-slide images of the lung sections were captured with the EVOS M7000 Images System (Thermo Fisher Scientific). Microscopic evaluation of pathological lung lesions was performed blindly by pathologists following a semiquantitative scoring system with the inclusion of three indicators (*49*): (i) alveolar septum thickening and consolidation; (ii) hemorrhage, exudation, pulmonary edema, and mucous; and (iii) recruitment and infiltration of inflammatory immune cells. For each hamster, three or four lobes of the lung were assessed independently, and the average score was calculated to indicate the overall pathological severity.

### Statistical analysis

The Mann-Whitney *U* test was used for the comparison between two independent samples. The uncorrected Kruskal-Wallis test, Dunnett’s Multiple Comparison test, or uncorrected Fisher’s LSD test was applied to analyze differences among more than two groups. A two-sided log-rank test was applied to compare the difference in survival. Statistical differences were considered to be significant for two-tailed *P* values of < 0.05. All statistical analyses were conducted in GraphPad Prism 8 software.

## Author contributions

H.Z., Q.Y., Y.G., N.S.X. conceptualized and designed the study. S.W., K.W.,Y.W., Y.Z., M.W., J.X. designed the clones, and produced and characterized the proteins. Y.W., L.Y., J.M., M.Z., C.L., W.G. performed the animal experiments. H.C., Z.L., H.Z., X.W., Y.W. performed the rhesus monkey experiments. Z.W., Y.H., D.C., M.L., H.L.G., J.G., H.X. performed the ELISA and neutralization assays. J.J.X., H.Y.G. performed ELISpot assays. J.M., P.C. performed the SARS-CoV-2 RNA quantification. M.N., Y.W. formulated the vaccines. Y.S., W.H. prepared mAbs. Q.Z. carried out the cryo-EM studies. Q.Y., Y.W., S.W., Q.Z., Z.H. wrote the paper. S.L., T.C., T.W., Y.C., Q.Z., J.Z., T.Z., H.Z., Y.G., N.S.X. revised the manuscript. All authors read and approved the final version of the manuscript.

## Data and materials availability

Structure coordinates are deposited in the Protein Data Bank under accession codes 7WP6 (STKF:36H6:83H7:85F7) and 7WP8 (STFK1628x:83H7:85F7:2B4). The corresponding EM density maps have been deposited in the Protein Data Bank under accession numbers EMD-32676 (STKF:36H6:83H7:85F7), EMD-32678 (STFK1628x:83H7:85F7:2B4). All data associated with this study are present in the paper or the Supplementary Materials. Reagents will be made available to the scientific community by contacting N.S.X. or Q.Y. and completing a materials transfer agreement.

---

**fig. S1.**
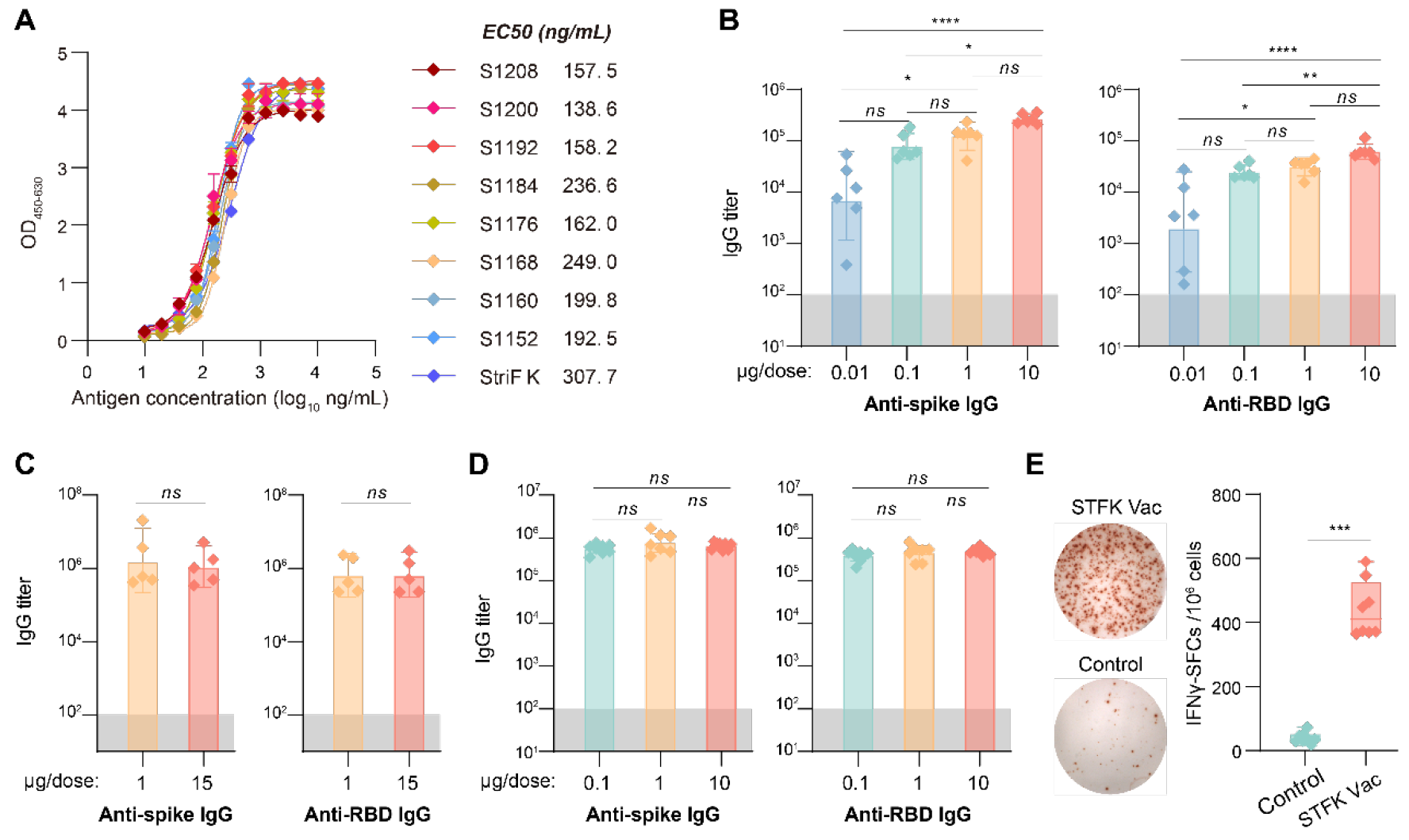
Evaluations for the recombinant STFK for *in vitro* binding with human rACE2 and *in vivo* immunogenicity. **(A)** ELISA-binding activities of recombinant spike proteins with human rACE2. **(B-D)** Anti-Spike and anti-RBD IgG titers in STFK-immunized **(B)** BALB/c mice, **(C)** rhesus monkeys, and **(D)** hamsters. **(E)** Spike-specific T cell response elicited by STFK-vaccination in C57BL/6 mice measured by ELISpot assays. Representative images (right panel) and the counts of IFN-γ spot-forming cells (left panel) were shown. Data in (B-D) were plotted as the geometric mean with SD. Data in (E) were shown as box and whisker plots; median, first quartile, third quartile, minimum value, and maximum value were plotted. Dark shadows in (B-D) indicate the LOD. Uncorrected Kruskal-Wallis test (B, D) or Mann-Whitney *U* test (C, E) were used for intergroup statistical comparisons. Asterisks indicate statistical significance (\*\*\*\**P* < 0.0001; \*\*\**P* < 0.001; \*\**P* < 0.01; \**P* < 0.05; ns, not significant).

**fig. S2.**
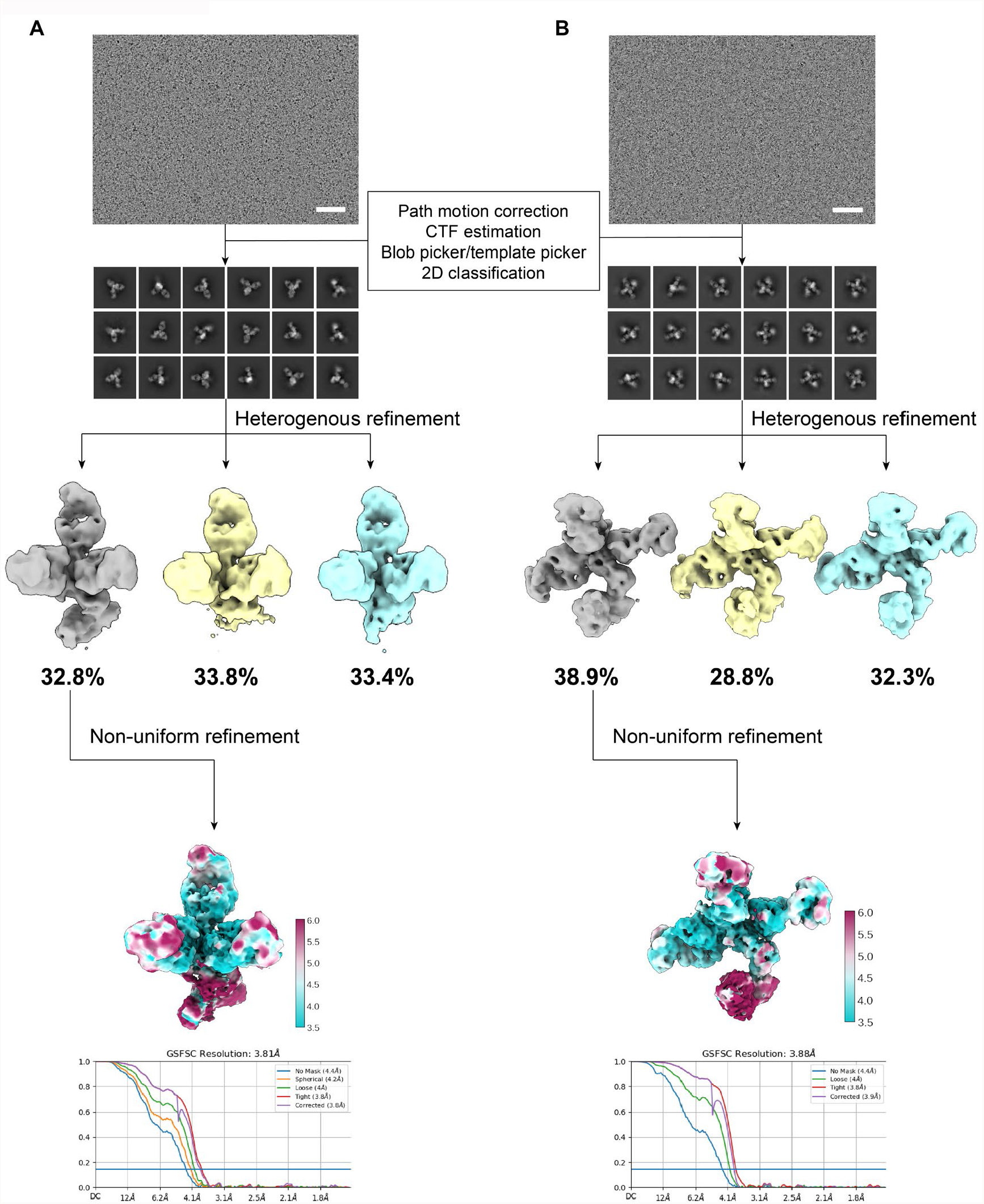
Flowchats of cryo-EM images processing and 3D reconstructions of STFK:36H6:83H7:85F7 (A) and STFK1628x:83H7:85F7:2B4 (B). Fourier shell correlation (FSC) curves and local resolution analysis of 3D and reconstructions are shown, scale bar=50 nm.

**fig. S3.**
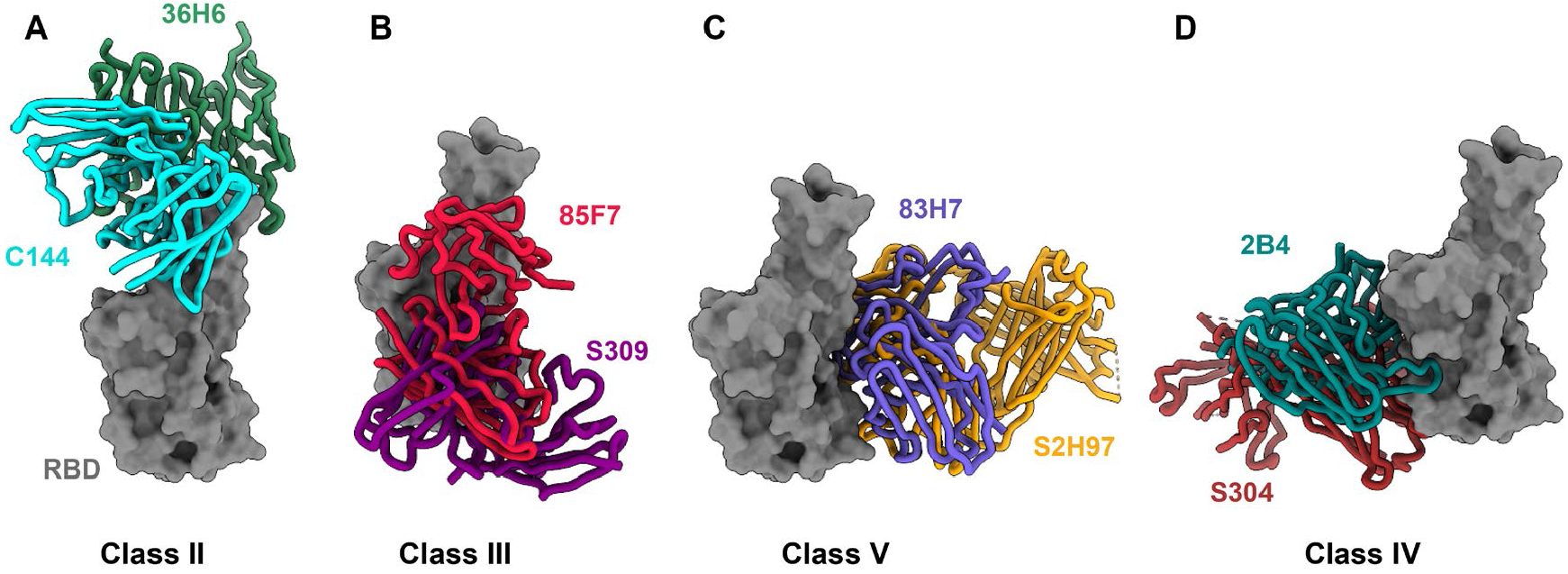
Comparison and classificatlion of nAbs by their binding epitopes and modes. 36H6 (A), 85F7 (B), 83H7 (C), and 2B4 (D) were grouped into Class II, III, IV, and V nAbs, and their binding modes are similar to reported nAbs C144 (Class II, pdb no. 7K90), S309 (Class III, pdb no. 7R6W), S2H97 (Class V, pdb no. 7M7W) and S304 (Class IV, pdb no. 7R6X), respectively.

**fig. S4.**
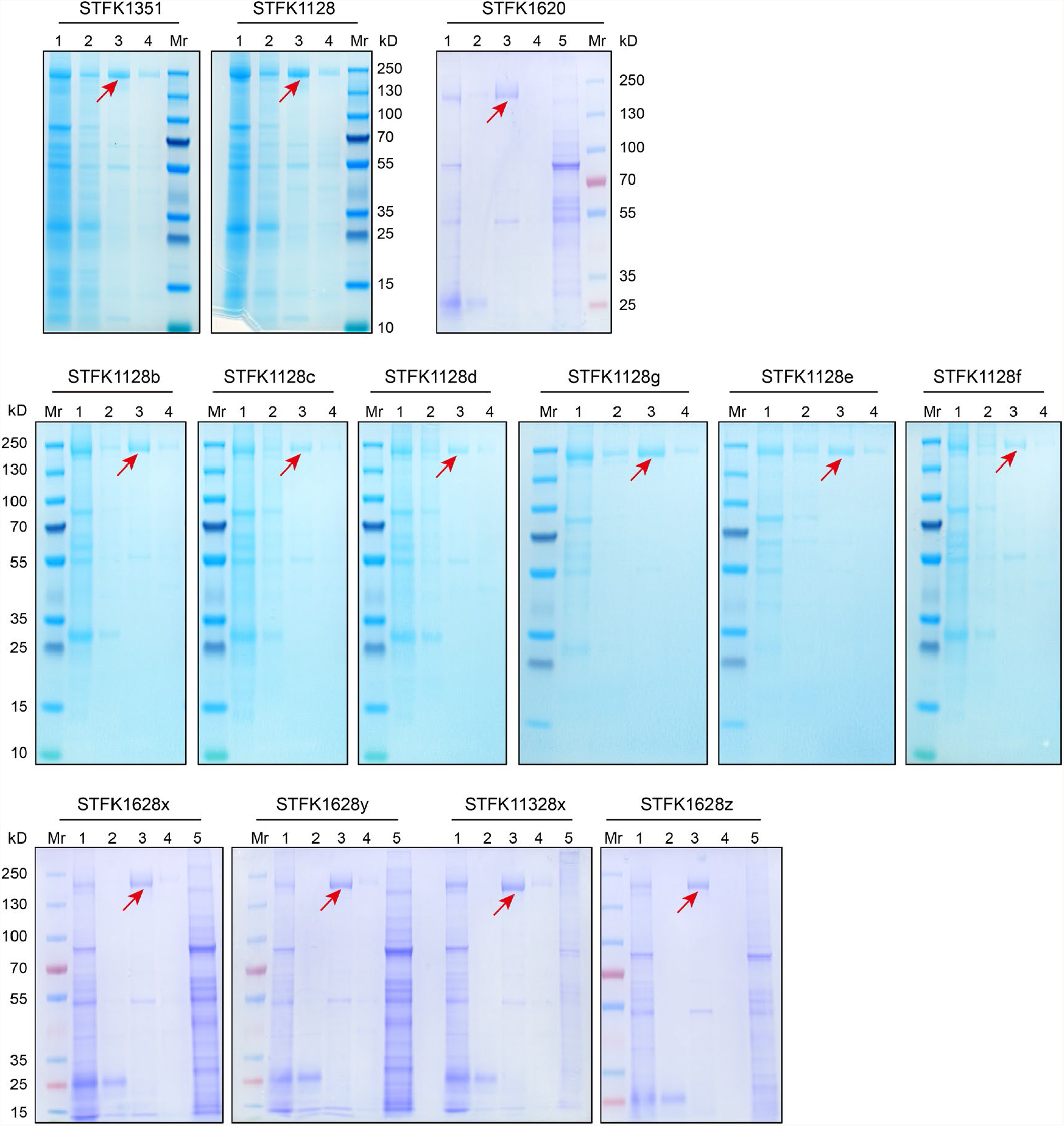
SDS-PAGE analyses for engineered STFK variants. Mr, protein ladder; lane 1, supernatants of transfected cells; lane 2, flow-through fraction from the Q-FF column; lane 3-4, the eluate fractions with buffer containing 100 mM NaCl; lane 5, eluate fraction with buffer containing 2 M NaCl. The red arrow indicates the target protein band.

**fig. S5.**
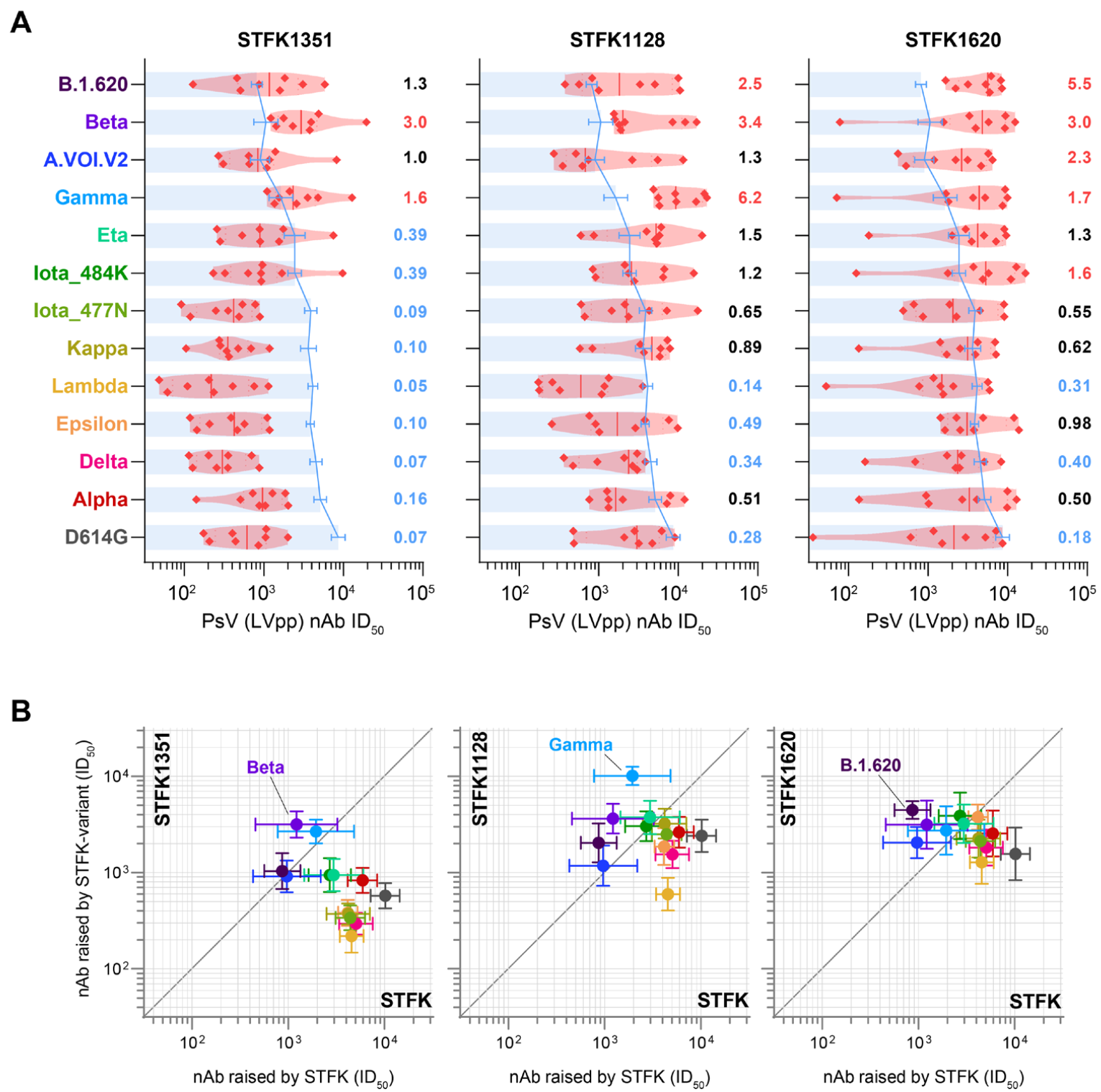
Neutralizing antibody responses elicited by STFK1351, STFK1128, and STFK1620 in hamsters. **(A)** The nAb titers of sera from hamsters (n=8) receiving vaccination of STFK1351, STFK1128, and STFK1620 to neutralize lentiviral-pseudotyped SARS-CoV-2 variants. The blue lines (bars) indicate the nAb GMTs (±SD) induced by the prototypic STFK vaccine against the corresponding variants. The numbers on the right represent the GMT fold-changes of nAb titers elicited by STFK variants to the prototypic STFK. The fold-changes were colored according to the values: <0.5 was in blue, 0.5-1.5 was in black, and >1.5 was in red. **(B)** The scatter plots compare the cross-neutralizing activities of nAbs raised by STFK variants (Y-axis) and prototypic STFK (X-axis). Data were plotted as the geometric mean with SEM. The diagonal line was Y=X.

**fig. S6.**
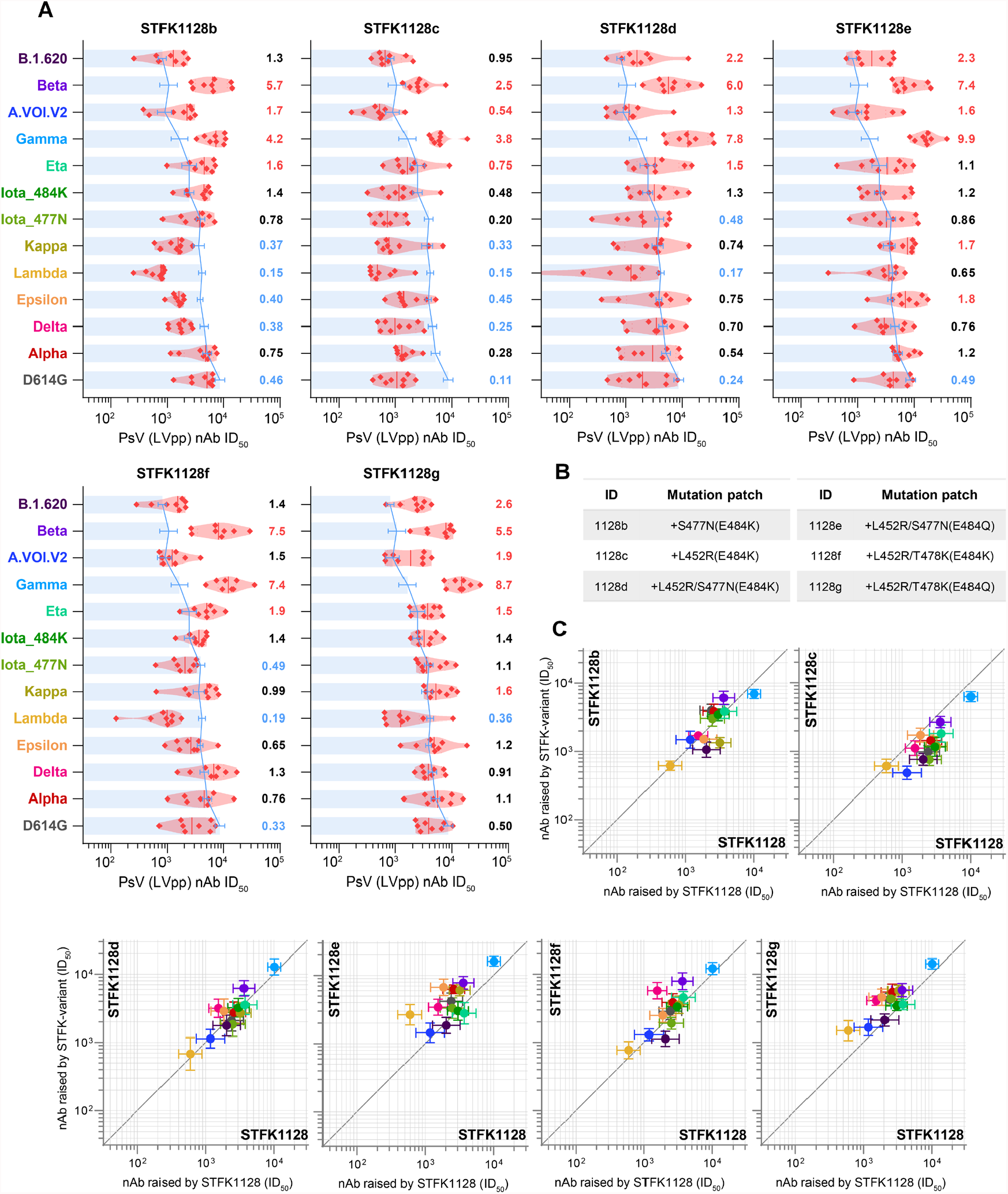
Neutralizing antibody responses elicited by STFK1128 derivates in hamsters. **(A)** The nAb titers of sera from hamsters (n=8) receiving vaccination of six STFK1128 derivates **(B)** to neutralize lentiviral-pseudotyped SARS-CoV-2 variants. The blue lines (bars) indicate the nAb GMTs (±SD) induced by the prototypic STFK vaccine against the corresponding variants. The numbers on the right represent the GMT fold-changes of nAb titers elicited by STFK variants to the prototypic STFK. The fold-changes were colored according to the values: <0.5 was in blue, 0.5-1.5 was in black, and >1.5 was in red. **(B)** Additional mutation patches in STFK1128 derivates compared to its parental construct. **(C)** The scatter plots show the comparison of nAbs against 13 variants raised by STFK1128 derivates (Y-axis) and their parental STFK1128 (X-axis). Data were plotted as the geometric mean with SEM. The diagonal line was Y=X.

**fig. S7.**
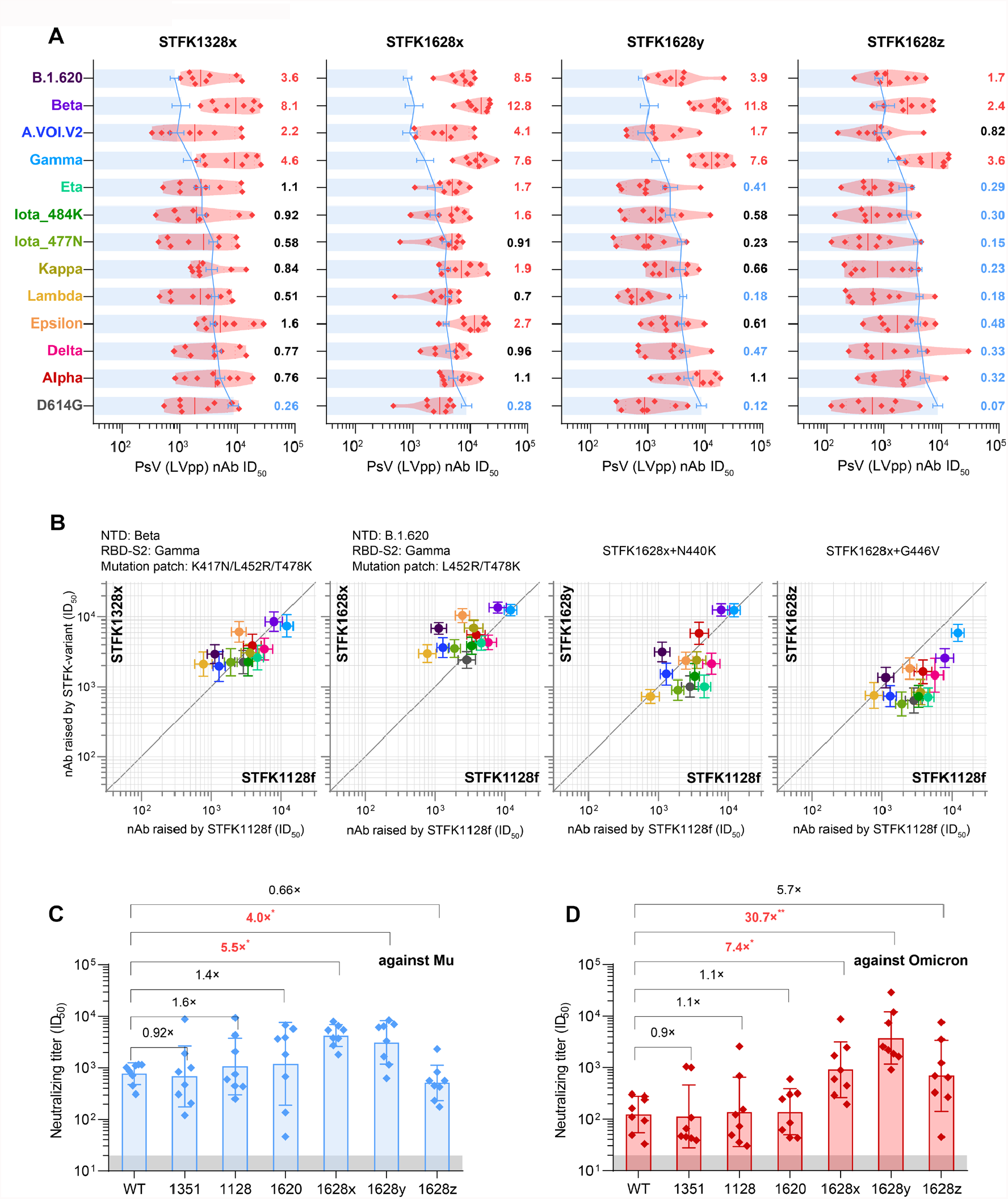
Neutralizing antibody responses elicited by inter-lineage chimeric STFK variants in hamsters. **(A)** The nAb titers of sera from hamsters (n=8) receiving vaccination of four STFK1128f-derived chimeric STFK variants to neutralize lentiviral-pseudotyped SARS-CoV-2 variants. The blue lines (bars) indicate the nAb GMTs (±SD) induced by the prototypic STFK vaccine against the corresponding variants. The numbers on the right represent the GMT fold-changes of nAb titers elicited by the chimeric STFK variants to the prototypic STFK. The fold-changes were colored according to the values: <0.5 was in blue, 0.5-1.5 was in black, and >1.5 was in red. **(B)** The scatter plots compare the cross-neutralizing activities of nAbs raised by chimeric STFK variants (Y-axis) and their parental STFK1128f (X-axis). Data were plotted as the geometric mean with SEM. The diagonal line was Y=X. A detailed information summary of each chimeric variant was shown on the top of the panels. **(C, D)** The nAb titers against the newly emerged variants of Mu **(C)** and Omicron **(D)** were elicited by the engineered chimeric STFK variants in comparison to STFK antigens based on naturally occurring variants. The numbers on the top of panels represent the relative nAb GMT changes induced by the STFK variants to the prototypic STFK. Dark shadows indicate the LOD. Uncorrected Kruskal-Wallis tests were used for statistical comparison. Asterisks indicate statistical significance (\*\*\*\**P* < 0.0001; \*\*\**P* < 0.001; \*\**P* < 0.01; \**P* < 0.05; ns, not significant).

**fig. S8.**
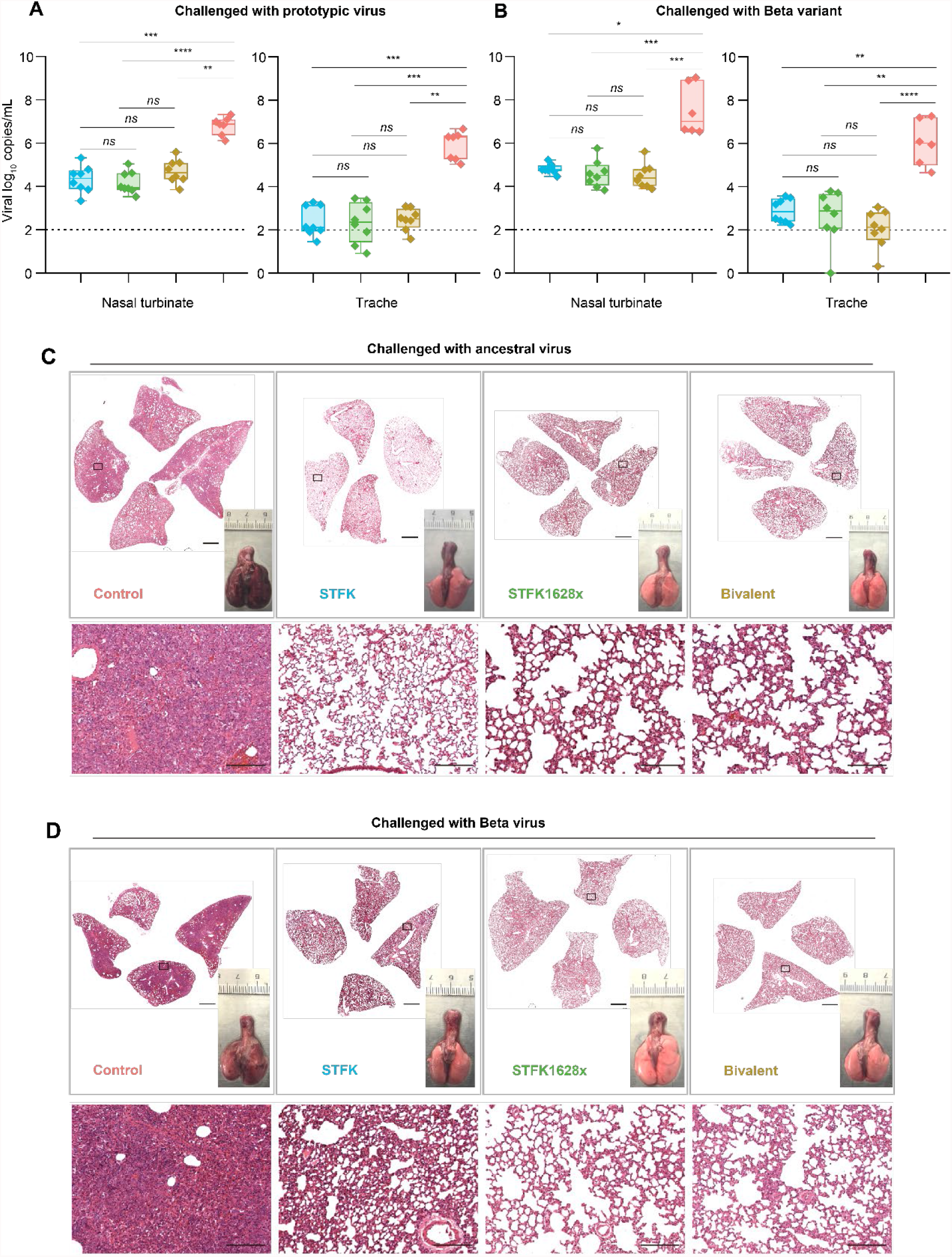
Tissue analyses for hamsters intranasally challenged with SARS-CoV-2. Animals were identical to that shown in Fig. 3. **(A-B)** Viral RNA levels in tissues of nasal turbinate (left panel) and trachea (right panel) collected from hamsters challenged with ancestral SARS-CoV-2 **(A)** or Beta variant **(B). (C-D)** Representative H&E-stained lung sections were collected from ancestral SARS-CoV-2 **(C)** or Beta variant **(D)** challenged hamsters. Views of the whole lung lobes (four independent sections) and the gross observations of lung tissues were presented in the top panel (scale bars, 2 mm), areas in the black box were enlarged in the bottom panel (scale bars, 200 μm). Uncorrected Kruskal-Wallis tests were used for intergroup statistical comparison. Asterisks indicate statistical significance (\*\*\*\**P* < 0.0001; \*\*\**P* < 0.001; \*\**P* < 0.01; \**P* < 0.05; ns, not significant).

**Table S1.** Cryo-EM data collection, refinement and validation statistics of three-antibody immune-complexes.

|  | STFK:36H6:83H7:85F7 | STFK1628x:83H7:85F7:2B4 |
| --- | --- | --- |
| **Data collection and processing** |  |  |
| Microscope | FEI TF30 | FEI TF30 |
| Camera | K3 | K3 |
| Magnification | 39,000 | 39,000 |
| Voltage (kV) | 300 | 300 |
| Electron exposure (e-/Å<sup>2</sup>) | 60 | 60 |
| Defocus range (μm) | 1.2-3.5 | 1.0-3.0 |
| Pixel size (Å) | 0.778 | 0.778 |
| Micrographs (total) | 3,479 | 4,191 |
| Micrographs (used) | 2,576 | 3,773 |
| Total particle | 1,146,590 | 1,684,307 |
| Final particle images (no.) | 162,177 | 115,589 |
| Symmetry imposed | C1 | C1 |
| Map resolution (Å) | 3.81 | 3.88 |
| FSC threshold | 0.143 | 0.143 |
| Map sharpening B factor (Å<sup>2</sup>) | -142.4 | -119.8 |
| **Validation** |  |  |
| MolProbity score | 1.96 | 1.99 |
| Clashscore | 7.71 | 8.64 |
| Poor rotamers (%) | 0.69 | 0.00 |
| RMS (bonds) | 0.0113 | 0.0087 |
| RMS (angles) | 1.36 | 1.32 |
| Ramachadran plot |  |  |
| Favored (%) | 90.44 | 90.91 |
| Allowed (%) | 9.44 | 8.97 |
| Disallowed (%) | 0.12 | 0.12 |

